# Evidence for *in vivo* mRNA Transport Between Mammalian Cells

**DOI:** 10.1101/2024.10.11.617465

**Authors:** Andrey Shubin, Songlei Liu, George Church, Craig Hunter

## Abstract

The prevalence and significance of intercellular mRNA transport remains unknown. Direct detection of mRNA transfer between cells within an organism is challenging due to technical limitations associated with transgene encoded molecular labels and cell sorting techniques. In this study we analyzed human-to-mouse xenograft single-cell RNA sequencing data to identify mouse transcripts in recovered human cells. The murine transcriptome analysis implicates macrophages as a frequent mRNA donor cell type. We then developed an *in vitro* system using mouse RAW264.7 macrophages and human HeLa and HEK293 cells and confirmed the transfer of Ftl1 mRNA from mouse to human cells. Overall, our study provides compelling *in vivo* evidence for prevalent intercellular mRNA transfer in human-to-mouse xenograft models.

## Introduction

Intercellular communication in multicellular organisms plays a vital role in development and physiology. Multiple pathways of juxtracrine, paracrine and endocrine communication are mediated by diverse biological agents including RNA-carrying extracellular vesicles^1^. Finding these extracellular entities in many biological fluids (blood, cerebrospinal fluid, saliva, and urine) has led to the tantalizing possibility that extracellular RNA plays an active role in intercellular communication^2,3^. Studies in plants and nematodes support this hypothesis, demonstrating that the intercellular transfer of mRNA, small interfering RNA, and double-stranded RNA enable non-autonomous gene expression and systemic gene silencing^4–12^. While studies in mammalian cells support RNA transfer as a form of intercellular communication, the evidence is indirect or confined to very specific cases. This includes the detection of products of translation from the candidate transferred transcripts (mRNA)^13,14^, changes in gene expression levels in recipient cells (miRNA)^15^, or restoration of RNA-specific functions in knockout recipient cells (snoRNA)^16^. However, these functional readouts are indirect and do not rule out the influence of confounding factors, including the co-transfer of proteins or other biological agents that could mediate the observed effects attributed to RNA transfer, as well as the unknown potential response of recipient cells to experimental stimuli^2^. Furthermore, the functional consequences attributed to RNA transfer are often demonstrated in *in vitro* systems that do not reflect the *in vivo* context, for example, exposure to non-physiological levels of exogenous RNA^17,18^. Such approaches blur the distinction between what can occur *in vitro* from what happens *in vivo*.

The objective of this study was to obtain direct evidence of intercellular mRNA transport by identifying transcripts that are synthesized in donor cells and transferred into acceptor cells. The fundamental limitation in studying mRNA transfer is the inability to distinguish between donor and acceptor cell mRNA. In *in vitro* models, labelling mRNA from donor cells can be accomplished with modified nucleotides. However, it is challenging to discriminate between the transfer of labeled mRNA molecules from the transfer of free modified nucleotides that acceptor cells can utilize for their own mRNA synthesis. A more effective approach involves using donor and acceptor cells derived from different species. In this case, once mRNA from the donor cell is transferred to the acceptor cell, it can be distinguished by its species-specific nucleotide sequence, enabling a clear distinction between donor cell transferred transcripts and acceptor cell endogenous mRNA. The simplest *in vitro* model for studying mRNA transfer involves co-culturing cells from different species followed by single cell RNA-seq (scRNA-seq) to detect transferred transcripts. However, our understanding of intercellular mRNA transfer is still limited, leaving key aspects of the model undefined: the cell types most likely to serve as mRNA senders and receivers, the scale and intensity of the transfer, whether it occurs continuously or requires specific triggers, and whether all transcripts or only specific ones are transferred. This lack of critical information hinders the rational design of *in vitro* models and does not allow any benchmark to discriminate between a natural process from an unintended consequence of the experimental setup.

Furthermore, *in vitro* cell culture manipulations associated with direct co-culturing of adherent cells used in some studies of mRNA transfer^19^ can inadvertently result in mRNA transfer, making it extremely challenging to reliably and reproducibly control for these confounding factors^20,21^. Therefore, studying mRNA intercellular transfer based only on *in vitro* models with direct co-culture of cells may give an inaccurate representation of the process’s feasibility under physiological conditions.

To overcome these limitations, we first gathered primary data on mRNA transfer from *in vivo* animal models, where all the necessary parameters for this process are naturally in place, and then used these findings to rationally design *in vitro* confirmatory experiments. The most straightforward *in vivo* model to study mRNA transfer is to co-culture human cells engrafted into immunodeficient host mouse. mRNA transfer between mouse and human cells is then detected by re-isolating the human cells and immediately performing scRNA-Seq on the human cells. Most transcripts will be human, confirming the identity of the recipient cell, while any mouse transcripts are candidate host-derived transported mRNAs. This model represents a specific case within the broader context of human-to-mouse xenograft models, which are widely used in oncology and immunology research^22^. In our study, we identified three previously published xenograft studies that aligned with the design of our *in vivo* model to investigate mRNA transfer^23–25^ (Table 1). By re-analyzing scRNA-Seq data from these studies, we were able to identify transplanted human cells that showed evidence of murine mRNA transfer, detect specific murine transcripts associated with human cells, and identify murine macrophages as likely mRNA donor cells. Based on these findings, we designed an *in vitro* model and successfully recapitulated transfer of mouse transcript Ftl1 from immortalized mouse macrophage cells (RAW264.7) to HEK293 and HeLa human cells.

**Table 1.** Human-to-mouse xenograft studies and a barnyard control study selected for analysis of mouse-to-human mRNA transfer.

|  | GSE73122 | GSE96981 | GSE83142 | GSE63473<br>(contamination control) |
| --- | --- | --- | --- | --- |
| Xenoransplanted cells/tissues | Paired primary (**pRCC**) and metastatic (**mRCC**) human renal tumors. | Human iPS-derived liver buds (**Lb**). | Human patient-derived acute lymphoblastic leukemia cells (**ALL**). | **Barnyard experiment:** mixed human HEK293 and mouse 3T3 cells. |
| Xenotransplantation site | Subrenal capsule | Cranial window | Introduced via tail vein, recovered from bone marrow | - |
| Sequenced cell populations | 1) pRCC<br>2) mRCC | 1) Lb, day 3 p-t.<br>2) Lb, day 10 p-t.<br>3) Lb, day 15 p-t. | 1) **Non-LRC**, proliferative ALL, 15 days p-t.<br>2) **LRC**, dormant ALL, 15 days p-t.<br>3) **PDXt**, ALL treated with combination of anti-cancer drugs.<br>4) **PDXnt**, ALL not treated with anti-cancer drugs. | Sorted HEK293 cells immediately processed for single cell RNA-seq. |

In summary, our *in vivo* data supports mRNA transfer in human-to-mouse xenograft models, while our *in vitro* data provides direct evidence for intercellular mRNA transfer from macrophage cells.

## Results

To determine the prevalence and extent of intercellular mRNA transport we re-analyzed single-cell RNA-seq (scRNA-seq) data derived from human-to-mouse xenograft studies (Fig. 1). This approach identified candidates for intercellular mRNA transfer without the requirement for nucleic acid or cell labeling. Importantly, scRNA-seq analysis allowed us to explore the entire murine transcriptome for mobile mRNAs. To improve our ability to detect rare and potentially unstable mobile mRNAs, we selected studies in which potential recipient cells recovered from xenograft animals were processed for scRNA-seq without interim culturing. The two most widely used platforms for high-throughput scRNA-seq are 10x Genomics and C1 Fluidigm. 10x Genomics offers the advantage of simultaneously profiling the transcripts of tens of thousands of cells, enabling the detection of rare RNA transfer events. However, the reduced sequencing depth and inability to cover the entire transcript characteristic of this platform may limit the ability to detect transferred RNA molecules. In contrast, the microfluidic chip-based C1 Fluidigm RNA-seq platform, although limited to 96 cells per chip, provides several million reads per cell. If RNA transfer is biologically significant, it is likely to occur frequently enough to be captured among a few dozen cells. Moreover, the high read depth offered by the C1 Fluidigm platform ensures sufficient sensitivity to capture multiple instances of transported RNAs. We identified three studies that utilized the Fluidigm platform, each involving several populations of cells engrafted into different tissues and incubated in host animals for varying durations and under diverse conditions (Table 1)^23–25^.

**Figure 1.**
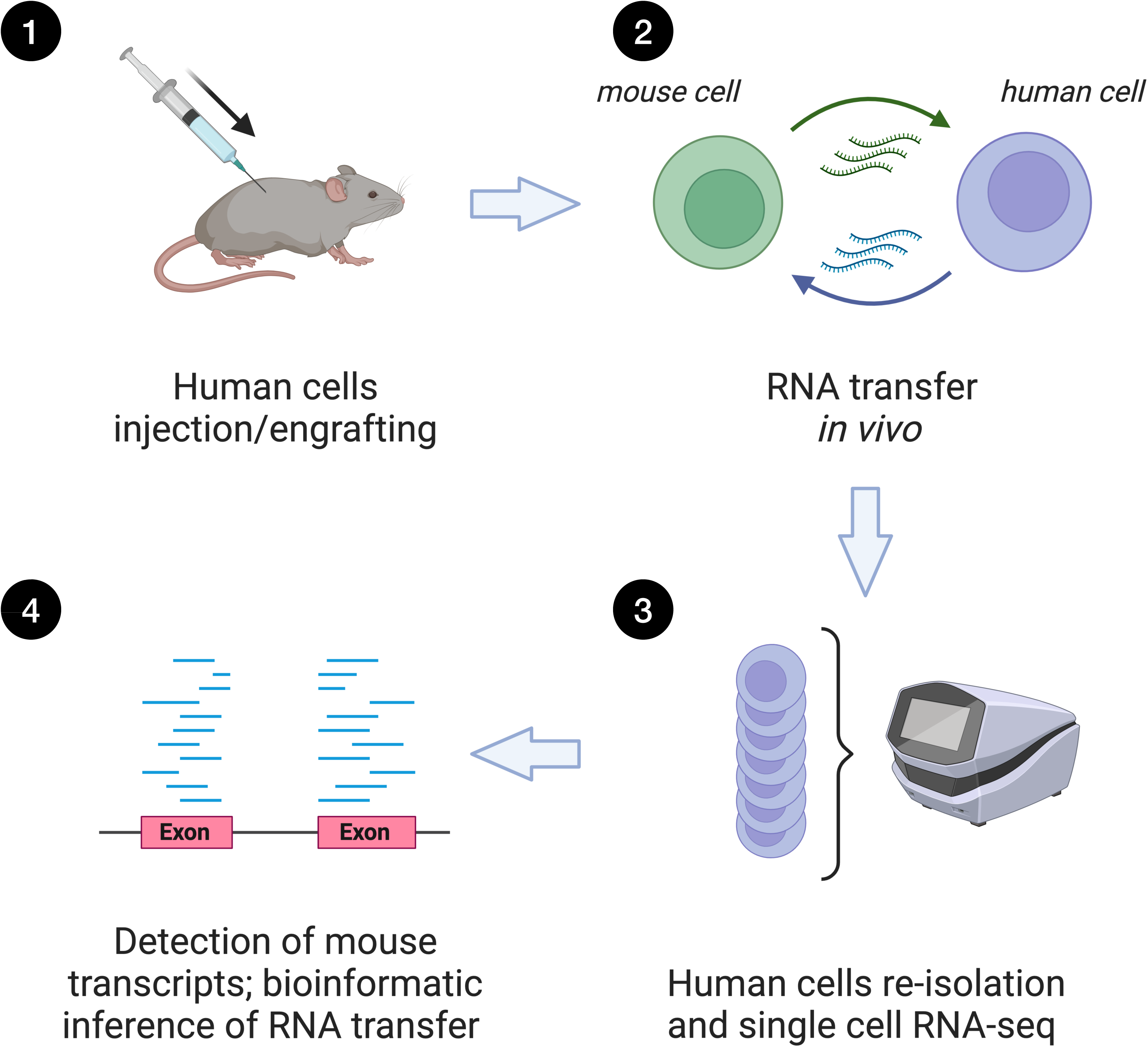
Design of the studies selected to detect RNA transfer. ***1.*** Engrafting of human cells into host mice. ***2.*** Culturing of the grafted cells within the host animal. ***3.*** Re-isolation of the grafted cells followed by immediate processing for RNA-seq analysis. ***4.*** Bioinformatic separation of human and mouse-specific reads to identify mouse transcripts associated with engrafted human cells.

Two challenges associated with the use of scRNA-seq are sequencing errors, which can introduce uncertainties regarding the species identity of specific reads, and the need to differentiate between intercellular RNA transport and inadvertent host-cell contamination that may occur during cell dissociation. To address the first challenge, we established a minimum mRNA detection threshold of four non-duplicated high-complexity mouse-specific reads per transcript. Additionally, to mitigate the risk of misalignment of mouse reads to the human genome, we incorporated a read filtration step using the *blastn* algorithm within our bioinformatics pipeline (Fig. S1; see Methods for details). The second challenge, distinguishing between contaminating host RNA and potentially transported RNA, is further complicated by the fact that the selected studies were not specifically designed to detect intercellular RNA transport. Since the selected datasets lack contamination controls, we identified a scRNA-seq specificity study that purposefully mixed mouse and human cells prior to selecting human cells for scRNA-seq on the C1 Fluidigm platform^26^ (Table 1). In this experiment the number of mouse-specific reads detected per human cell exhibited a wide range, varying from several dozen to nearly 1.5 thousand. The latter corresponds to nearly 100 mouse transcripts per cell (Fig. 2A). As anticipated, the detected mouse transcripts were dominated by highly abundant RNAs, including 18S rRNA, mitochondrial transcripts, and transcripts encoding ribosomal proteins (Fig. 3A). These results indicate that when donor and acceptor cells are individually cultured, dissociated from the substrate, mixed, and immediately processed using the C1 Fluidigm single-cell platform, a significant level of contamination with ambient RNA can occur. Therefore, we regarded the high detection frequency of each of these mouse transcript types in human cells as a distinct contamination signature.

**Figure 2.**
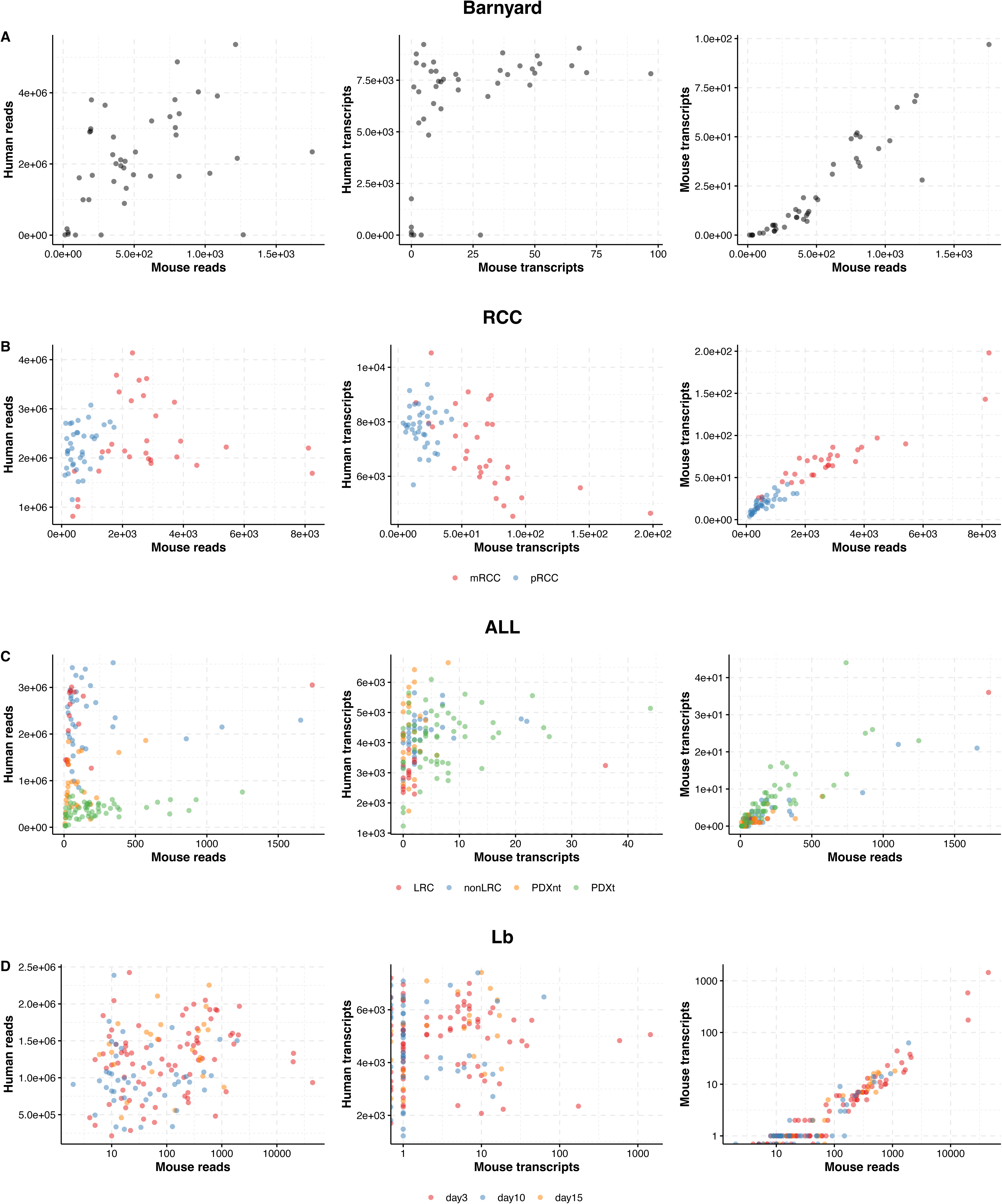
Distribution of re-isolated human cells by the number of mouse- and human-specific RNA-seq reads and corresponding transcripts. Only cells with ≥4 unique mouse-specific reads per identified mouse transcript are shown. Every dot corresponds to a single cell. Colors represent cell populations according to the legend. All abbreviations are used as in Table 1. Plots on figure D have log scale on x axis. For the full list of read counts and identified transcripts see Supplemental Table S1.

**Figure 3.**
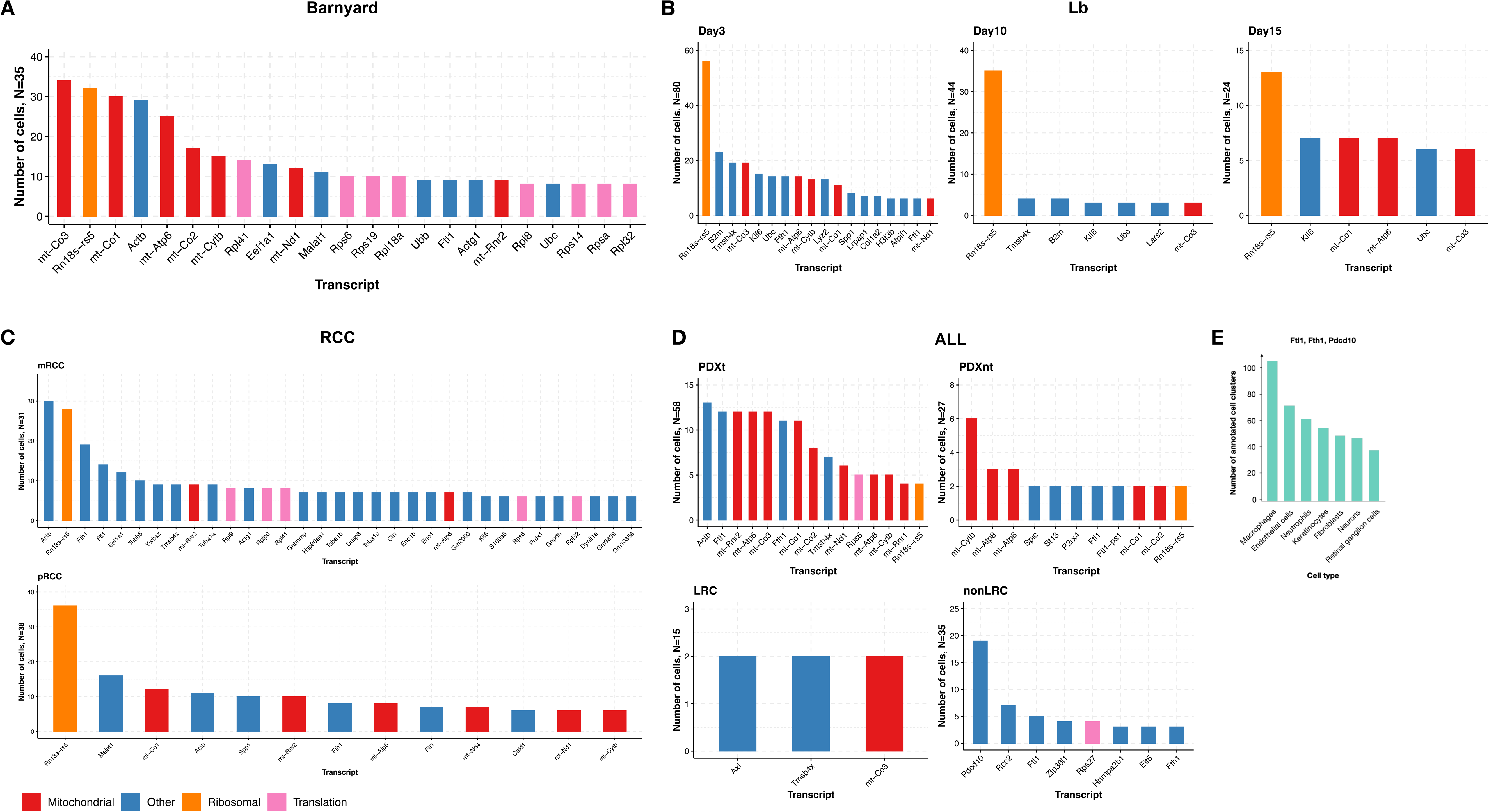
Identity and characteristics of most frequent mouse transcripts recovered from human cells. A-D, mouse transcripts most frequently detected in the re-isolated human cells from the re-analyzed studies. For the full list of identified mouse transcripts see Supplemental Table 2. A, The Barnyard study is a contamination positive control^26,41^. B, C, and D, are liver bud (LB), Renal carcinoma cancer (RCC) and Acute lymphatic Leukemia (ALL) studies respectively^23–25^. E, percentage of spliced transcript covered with sequencing reads for the mouse mRNA most frequently found in nonLRC cells from ALL study (panel D). F, cell types listed by the number of cell clusters with simultaneous expression of transcripts Pdcd10, Ftl1, and Fth1 from PanglaoDB database^27^.

The filtered and mapped mouse reads/transcripts obtained from the sequenced human cells in each of the three identified studies (Table 1; abbreviated as ALL (acute lymphoblastic leukemia), RCC (renal cell carcinoma), and Lb (liver bud organoids)) in general exhibited human to mouse reads ratio within the range found in the barnyard study. The relationship between the number of mouse reads and the number of mouse transcripts was roughly linear with no dependency between the numbers of human and mouse transcripts (Fig. 2 B-D). However, there was variability in the manifestation of contamination signatures in the individually prepared cell populations within each study (Fig. 3 A-D). Both the RCC and Lb studies displayed at least two of the three contamination signatures (presence of ribosomal RNA, and/or mitochondrial RNA, and/or mRNA encoding ribosomal proteins) within their respective individual cell populations (Fig. 3 B, C). In contrast, two of the four cell populations analyzed in the ALL study did not exhibit a strong contamination signal (Fig. 3 D). In this study^24^, patient derived ALL cells were labeled with the proliferation-dependent dye, which was used to distinguish between slowly growing cells that retain the dye (label-retaining cells, LRC) and cells that proliferate rapidly and fail to retain the dye (nonLRC). The labelled cells were injected into a host mouse via the tail vein and re-isolated from the bone marrow. Among the 15 sequenced slow growing cells, only two mouse nuclear transcripts (Axl, Tmsb4x) were found in only two cells, and 35 other mouse nuclear transcripts were found only in one cell (SRR3654862; Fig. S3 A). In contrast, many more actively proliferating cells contained multiple mouse transcripts (Fig. S3 B). Among them, Pdcd10, Rcc2, and Zfp36I1 exhibit consistently low expression levels in mouse bone marrow, yet they were detected in a similar or greater number of cells compared to the highly expressed Ftl1 and Fth1 transcripts. (Fig. 3 D, Supplemental Table 2).

The above analysis indicates that the proliferating cells may be enriched for transferred mouse RNAs relative to contaminating mouse RNAs. In the ALL study, the sequencing libraries were prepared with a SMARTer Ultra Low RNA Kit (Clontech) which allows efficient cDNA synthesis from full-length transcripts using an oligo-dT primer, therefore we were able to assess the integrity of the detected mouse transcripts. We hypothesized that RNAs released from mouse cells during single-cell isolation were less likely to remain intact compared to transferred RNAs. We identified many nonLRC human cells that contained mouse Pdcd10 with complete coverage of all coding exons, as well as several cells with complete coverage of coding sequence for mouse Flt1 and Fth1 transcripts (Fig. S2). This completeness of coverage was not observed for other detected transcripts in both LRC and nonLRC cells, where the coverage was only partial and biased towards the 3’ regions (see Supplemental archive 1 for *bam* files). Based on these findings, we selected Pdcd10, Ftl1, and Fth1 as candidates for intercellular mouse to human transported transcripts.

We proceeded to analyze the expression profiles of single-cell mouse transcriptomes using the Panglao Database^27^, which integrates multiple single-cell RNA-seq datasets from mice, to identify the most probable bone marrow cell types as mRNA donors.

Among the different bone marrow cell types, co-expression of Pdcd10, Ftl1, and Fth1 genes was most frequently found in macrophages (Fig. 3 F). We propose that macrophages likely serve as donors in mRNA intercellular transport *in vivo*.

We next designed an *in vitro* cell co-culturing system using immortalized mouse RAW264.7 macrophages as mRNA donor cells and human HEK293 or HeLa cells as mRNA acceptors. Detecting RNA transfer *in vitro* between co-cultured cells presents a particular challenge as the non-physiological environment can induce artifacts that resemble RNA transfer. It is known that when adherent cells initially spread on a substrate, there is an accompanying increase in exocytosis followed by a period of heightened endocytosis^20,21^. This enhanced exocytosis by donor cells can lead to non-physiologically high levels of extracellular vesicles carrying extracellular RNA, while increased endocytosis by acceptor cells can cause increased uptake of this extracellular material. These artifacts, either individually or in combination, can yield false-positive results. Additionally, debris from dead cells or cells damaged during trypsinization can serve as a source of extracellular RNA, which is practically impossible to completely exclude or control for when working with adherent cells. To minimize these known artifacts, we implemented several precautions. We avoided simultaneous or sequential plating of donor and acceptor cells in the same area for co-culturing. Instead, we cultured donor and acceptor cells in separate wells created by a removable insert. Only after the cells had fully settled and spread on the substrate was the insert removed, allowing the cells to grow and migrate towards each other; finally, recipient cells were assessed for intercellular RNA transfer only in the area where donor cells were not seeded. (Fig. 4A). This approach ensures that only migration-competent donor cells interact with acceptor cells.

**Figure 4.**
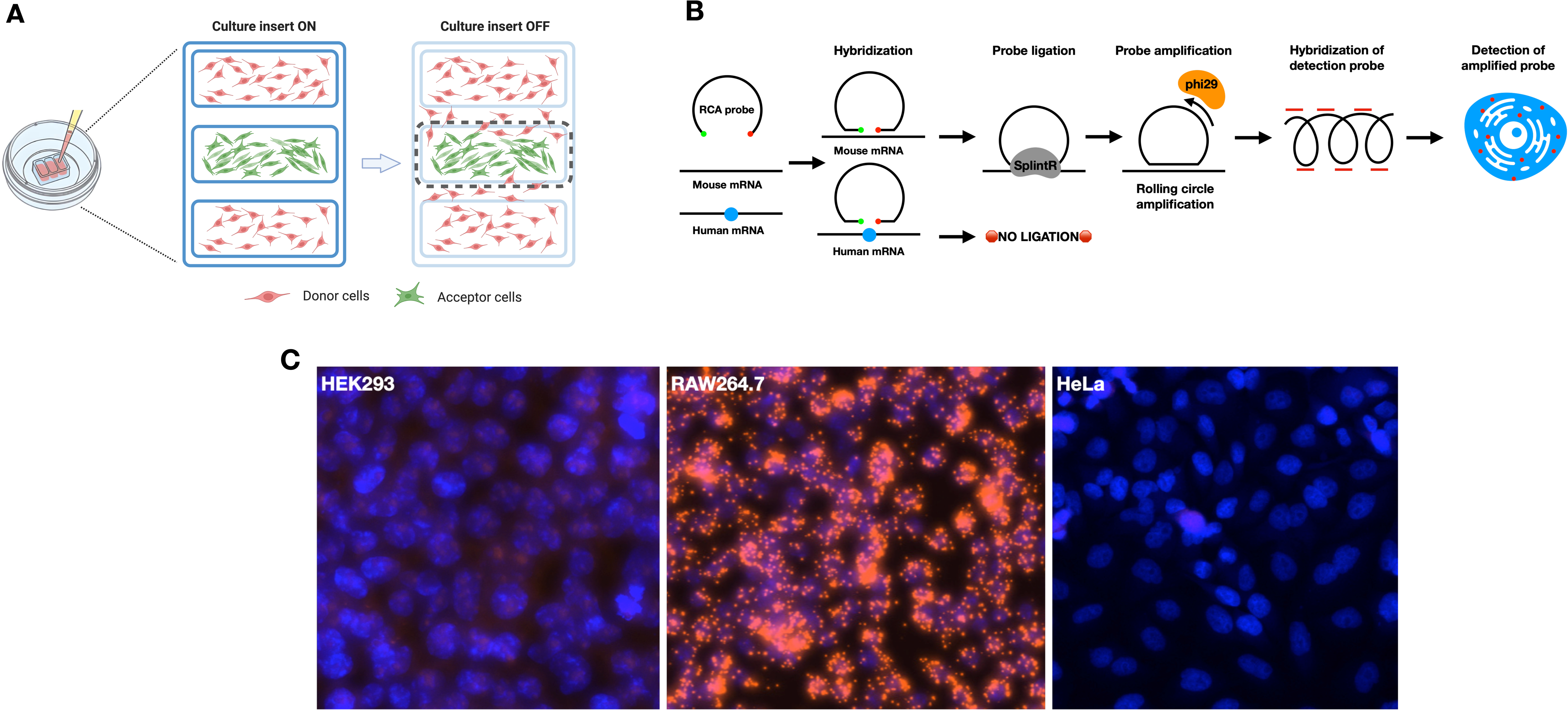
*In vitro* co-culturing system and highly specific detection of mouse transcripts. A. Set-up to grow spatially separate *in vitro* co-culturing of mRNA donor cells (RAW264.7) and mRNA acceptor cells (HeLa, HEK293). Cell near the margin (dashed line) were scored for transfer of donor cell mRNA in acceptor cells. B, layout of method of *in situ* mRNA-templated hybridization of ssDNA padlock probes. The species-specific sequence mismatch prevents probe ligation on human transcripts. The ligated probe is detected by rolling circle amplification followed by hybridization to probe sequences. C, assessment of specificity of discrimination between mouse (Ftl1) and human (FTL1) homological transcripts in the absense of co-culturing in cell types used as mRNA donors (RAW264.7) and acceptors (HeLa, HEK293) in *in vitro* system.

To detect mouse donor cell transcripts in single human receptor cells co-cultured with mouse cells, we employed *in situ* methods, which circumvent artifacts associated with trypsinization and cell sorting. While the hybridization conditions utilized in typical *in situ* hybridization are adequate for localizing and distinguishing different transcripts within a species, they are not sufficient to distinguish highly similar mouse and human homologs. To achieve both high sequence specificity and robust single molecule detection, we employed mRNA-templated hybridization and circularization of single-stranded DNA (ssDNA) padlock probes, followed by rolling circle amplification and detection using fluorescence in situ hybridization (FISH)^28^ (Fig. 4 B). Using this method, we were unable to detect Pdcd10 expression in RAW264.7 cells but observed low expression for Fth1 (data not shown) and high expression for Ftl1 (Fig. 4 C).

After 36-48 hours of co-culture, we observed active migration of RAW264.7 donor cells towards acceptor cells, while the latter exhibited little to no migration from their initially plated areas (Fig. 5 A, S4 A). Migrated RAW264.7 cells had many Ftl1 transcripts localized both in cell bodies and filopodia. In the regions where macrophages closely approached acceptor cells, we detected Ftl1 transcripts localized within the areas occupied by the acceptor cells (Fig. 5 B, S4 B). In these sites, we did not observe direct interactions of cell bodies or cell-to-cell contacts mediated by tunneling nanotubes through the transmitted light channel. However, we observed extracellular Ftl1-positive foci in the vicinity to the migrated RAW264.7 cells. The number of the extracellular Ftl1- positive foci decreased with the distance from the RAW264.7 cells. The source of the extracellular Ftl1 transcripts could be biological (Ftl1-containing extracellular vesicles secreted by donor cells) or technical artifact (movement of RCA colonies resulted from the cell treatment during the steps of RCA assay), or both; however, we could not resolve their origin within this system. The latter case – movement of RCA colonies - could result in association of the colonies with surface of the acceptor cells leading to false positive detection of RNA transfer. To determine whether this signal was associated with the cell surface or localized within the cytoplasm (i.e., inside the cell), we visualized cytoplasmic GFP in HEK293 cells. Our findings revealed recipient HEK293 cells with Ftl1 transcripts localized within their GFP volume (Fig. 5 C) excluding false positives due to movement of the RCA colonies and confirming intercellular transport of Ftl1 transcript.

**Figure 5.**
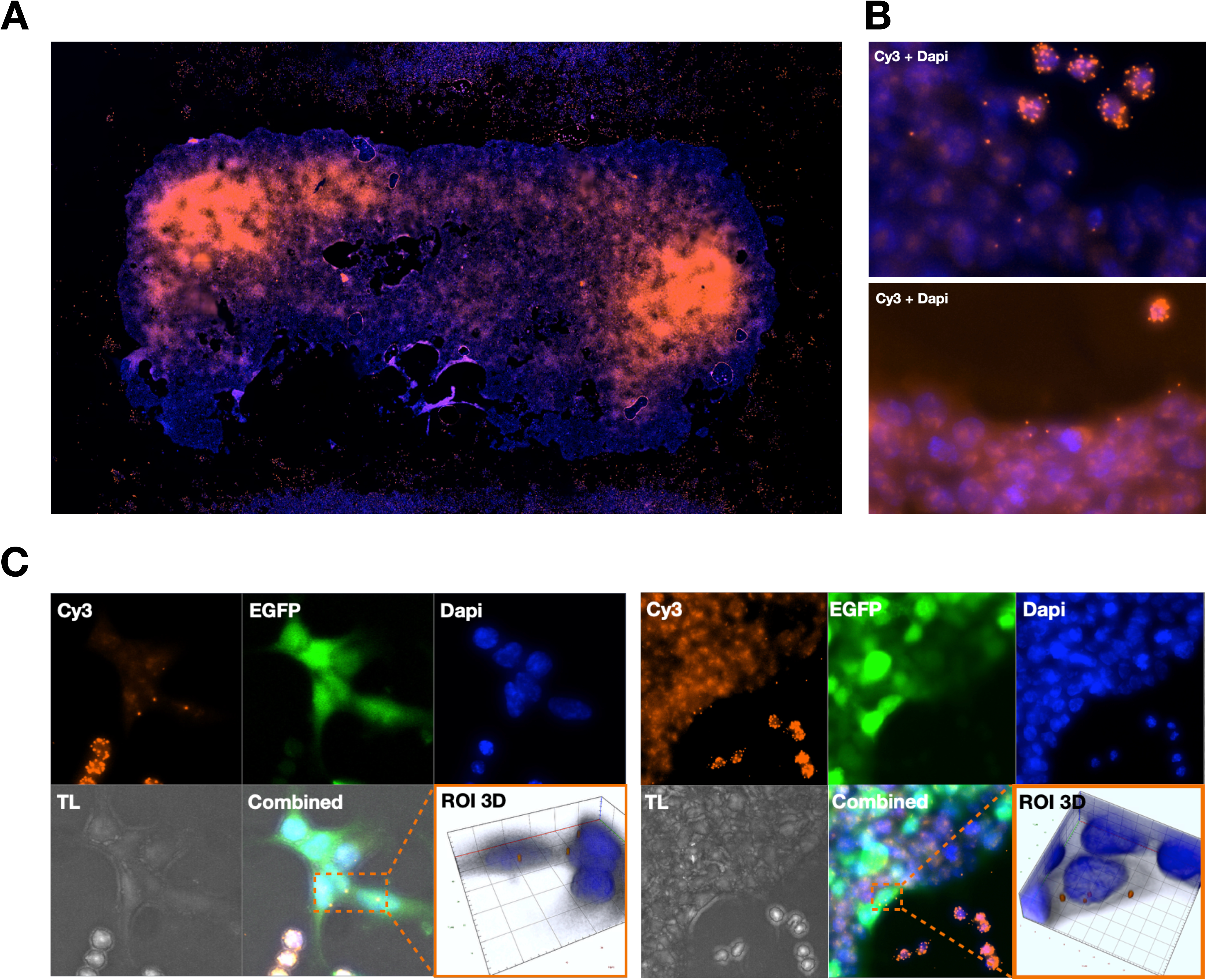
Detection of Ftl1 transfer in *in vitro* co-culturing system with HEK293- GFP cells as acceptors. A, panoramic view (maximum intensity projection) of the area occupied with seeded HEK293-GFP acceptor cells and sparse donor RAW264.7 cells that have migrated near the acceptor cells. B, representative examples from A of Ftl1 transcripts (labelled with Cy3 probes, shown in orange) localized within donor cells (many dots) and near or within the acceptor cells (sparse dots); DAPI staining is used for genomic DNA. C, examples of acceptor HEK293-GFP cells with Ftl1 transcript localized within the volume of visualized cytoplasmic GFP. ROI 3D, a 3D representation of the area highlighted by the dotted rectangle in the image; Cy3, cyanine 3, was used to visualize FISH probes to Ftl1-specific padlock probe; GFP, green fluorescent protein; DAPI, 4’,6-diamidino-2-phenylindole, was used to visualize genomic DNA; TL, transmitted light.

## Discussion

The goal of this study was to obtain definitive evidence for intercellular mRNA transfer in mammals and to assess its *in vivo* prevalence. We reasoned that scRNA-seq from human-mouse xenograft models would provide the specificity and sensitivity lacking in other hypothesis driven studies. This approach also avoids *a priori* assumptions about potential mechanisms, scale, and the cell types that act as donor and recipient cells.

Unfortunately, experimental systems that require physical isolation of cells from the host will always include risk of contamination. Therefore, we used three criteria to minimize false positive detection due to contamination during the process of cell isolation; 1) requiring multiple unique reads per transcript per cell, 2) detection of full-length transcripts, and 3) rejection of the cell populations with prevailing presence of at least one contamination signature inferred from the independent barnyard experiment. We then used this information to design a rigorous *in vitro* experiment to confirm transfer of Pdcd10 transcripts from macrophage like RAW264.7 donor cells to HeLa or HEK acceptor cells.

Among the three scRNA-seq studies we analyzed it was only the acute lymphoblastic leukemia (ALL) study where non-adherent human cells were recovered from the host animal’s bone marrow that met our contamination criteria. Because the ALL cells are non-adherent it was possible to recover and sort the cells at 4°C without trypsinization, which likely contributed to both stabilization of the transferred transcripts and minimized contamination of the recovered human cells with mouse transcripts by inhibiting their endocytotic activity during the processing. In contrast, isolation of adherent cells, either in *in vivo* or *in vitro* co-culturing systems, requires trypsinization that includes a 37°C incubation step. Trypsinization also results in cell rupture and release of RNA from donor cells either free or enclosed in endosome and exosomes. Furthermore, incubation at 37°C is favorable for endocytosis that can lead to engulfment of free RNA and RNA-containing vesicles by acceptor cells. It is not possible to properly control for trypsinization-mediated effects, and further separation and purification of acceptor cells is not helpful in this situation. Notably, although dormant and proliferative ALL cells were isolated from the same cell suspension by FACS, the proliferating cells contained many more mouse transcripts. This difference provides further evidence against prevalent contamination in these samples and can be attributed to the different endocytotic activity of proliferating (nonLRC) and dormant (LRC) cells.

The top three candidate transcripts - Pdcd10, Ftl1, and Fth1 – that were found to retain intact coding sequence in nonLRC ALL cells (15/35 for Pdcd10, 5/35 cells for Ftl1, 2/18 cells for Fth1; Fig. S2) – are very different in their expression ranks in mouse bone marrow. Pdcd10 - the most frequently detected transcript in the human acceptor cells - has much lower overall expression rank in mouse bone marrow compared to Ftl1 and Fth1. Since our design favors detection of selective mRNA transfer, we speculate we may have detected a special case of mRNA transfer from bone marrow cells with high expression of Pdcd10 induced in the context of ALL development.

Transcriptome profiling implicates macrophage like cells as the mRNA donor. Macrophages are known for their associations with tumor microenvironment and ambivalent role in tumor progression^29^. Since metastatic tumor cells acquire certain macrophage qualities, it has been proposed that macrophages could act to reprogram tumor cells from resting to migratory phenotype through complete or partial cell fusions (reviewed in Manjunath et al.^30^). In the case of partial fusion, it is assumed that partial sharing of macrophage transcriptome is sufficient to reprogram a tumor cell. Direct evidence of transmitting of portions of the cytoplasm from macrophages to melanoma cells resulted in an increase of migratory activity of tumor cells in *Zebrafish* xenograft system^31^.

We designed an *in vitro* co-culturing system to minimize false positive mRNA transfer detection by plating of donor and acceptor cells into separate but adjacent areas and by significantly raising the transcript detection threshold to maximize species-specific transcript discrimination. As described in the introduction and results sections, co-culture conditions can mimic intercellular RNA transfer. While detection of transcripts with RCA probes offers remarkable specificity, this depends on sequence context and often requires fine tuning of each probe’s binding sites to provide good sensitivity. Using RCA to discriminate between highly homologous transcripts brings significant limitations to the available probe binding sites. We choose to maximize species specificity, thus necessarily compromising detection sensitivity. Therefore, it is highly likely that the frequency and abundance of intercellular transfer we detected is significantly underestimated.

While this analysis and experimental follow-up were not designed to investigate the mechanism of RNA transfer, we speculate that it highly likely involves extracellular protein- or lipid-encapsulated RNAs, as we observed extracellular signal foci around RAW264.7 cells.

Furthermore, while we have no evidence that the transferred mRNAs are accessible to the translational machinery, it is noteworthy that reads originating from full-length transcripts were detected, leaving open the possibility of non-autonomy of genetic information. Non-cell autonomous RNA function has been proposed in plant root development^32^ and soma to germline transcript transfer has been detected in C. elegans^33^. However, the potential functional consequence for non-autonomous effects is perhaps greatest in non-human to human transplant patients.

In conclusion, considering the numerous technical and biological limitations of *in vitro* co-culturing and *in vivo* xenograft models, we suggest the future studies of intercellular RNA transfer operate on chimeric animals that are designed to have enough genetic heterogeneity but no or mitigated immune conflict^34^. Alternatively, analysis of recovered non-human tissues and organs from transplant patients or analysis of human cells from deceased transplant patients may provide evidence of RNA transfer. The development of sensitive *in situ* sequencing approaches, which would negate the requirement for and complications from cell isolation offers the best forward-looking prospects to evaluate the prevalence and functional consequence of intercellular RNA transfer.

## Methods

### Datasets and transcriptome selection criteria

Re-analyzed datasets are listed in the Table 1. Only single cell transcriptomes were selected for the analysis. Transcriptomes with any number of reads aligned to any of mouse hemoglobin genes were considered contaminated and discarded from the analysis.

### ScRNA-seq data re-analysis

RNA-seq data from the selected studies (Table 1) were re-analyzed from the level of *fastq* files (Fig. S1). As the goal of the analysis was only to identify mouse-specific reads, the read pairs in the paired-read data sets were split and reads were treated independently from their mates. The reads were initially aligned to GRCh38 human genome using *bowtie* aligner^35^ with no mismatches allowed (v=0). After that, unaligned reads were selected using *samtools*^36^, filtered for complexity by removing reads missing any of nucleotide types, and re-aligned to GRCm39 mouse genome (GCA_000001635.9) masked for all types of pseudogenes (annotation by Ensemble 110). Reads aligned to the mouse genome were deduplicated with *Rsubread* R package^37^ and additionally aligned to mouse and human genomes using *blastn* algorithm (ncbi-blast/2.5.0)^38,39^. Reads aligned only to mouse genome were considered mouse-specific. Both human and mouse-specific reads were counted using *HTSeq-count*^40^ in non-strand specific way (*stranded=no*). Further bioinformatics analysis was conducted using custom R scripts available upon request.

### Co-culturing of human and mouse cells in vitro

RAW264.7, HEK293-GFP, and HeLa cells were cultivated in DMEM culture medium (Gibco) supplemented with 10% FBS (Gibco) at 37C in 5% CO2 atmosphere. For the mRNA transfer *in vitro* assay, HEK293-GFP and HeLa cells were trypsinized, RAW264.7 cells were scraped off the substrate, washed several times in cold DMEM, and resuspended in full culture media at a density of 200-300 cells in 100 uL of full media. Cells were loaded into the wells of the Culture-Insert 3 Well (Ibidi, cat# 80366) mounted on a 18 mm poly-L-lysine-coated coverslip (neuVitro, cat# GG-18-1.5-pll) – RAW264.7 into the top and bottom cells, and HEK293-GFP or HeLa into the middle well. Cells were cultured for 36-48 h till ∼80% confluency followed by removal of the insert, transfer of the coverslip into 35mm culture Petri dish with 2mL of full culture media. Co-culturing lasted for 36-48 h until RAW264.7 cells migrate closely to the cells in the area of the middle well.

### In situ mRNA-templated hybridization

The assay was based on the protocol described by Liu et al^28^. Cells were twice rinsed in 2 mL of PBS (Corning) and fixed in 4% solution of paraformaldehyde (Sigma-Aldrich) in 1xPBS solution for 15 mins at room temperature, rinsed with phosphate-buffered saline (PBS) (GE Life Sciences), permeabilized with 0.25% Triton X-100 (Sigma-Aldrich) for 20 min and rinsed with Hybridization Buffer (10% deionized formamide (Ambion) in 6x SSC buffer (Thermo Fisher). Padlock probes (listed in Table S3) were hybridized at 100 nM final concentration in Hybridization Buffer for 1 h at 37°C, followed by three 5 min washes with Hybridization Buffer to remove unhybridized probe. After that, sample was rinsed with SplintR ligase reaction buffer (New England Biolabs) and incubated in 1x SplintR Ligase Reaction Buffer containing 210 nM SplintR Ligase (New England Biolabs) for 1 h at room temperature. After that, sample was rinsed three times with Hybridization Buffer and once with 1x Phi29 DNA Polymerase Buffer, followed by incubation with 1 U/μl NxGen Phi29 DNA polymerase (Lucigen, cat# 302210-2), 250 μM dNTPs, 200 μg/ml BSA (New England Biolabs), 200 nM of RCA primers (Table S3) in 1× Phi29 DNA Polymerase Buffer for 2 h at 37°C followed by three 5-min washes with 2x SSC buffer. Finally, the samples were stained with 500 nM Cy3 or Cy5 labelled FISH probes (Table S3) in 6x SSC for 30 min and washed 3 times with 6x SSC to remove unbound fluorescent probes. Sample was rinsed in water (Thermo Fisher, AM9938) and immediately mounted on a slide using Prolong Diamond Antifade Mountant (Thermo Fisher, P36970) and left to cure overnight. The specimens obtained were analyzed on Axio Scan.Z1 microscope (Zeiss).

## Author contribution

AS, CH –conceived the study, designed experiments and wrote the manuscript.

AS – carried out experiments and bioinformatics analysis.

SL – designed RCA probes and provided expertise in *in situ* mRNA-templated hybridization assay.

GC - provided expertise in *in situ* mRNA-templated hybridization assay.

## Acknowledgments

We thank Dr. Gray Camp, Dr. Christoph Ziegenhain and Dr. Kyu-Tae Kim for responding to our requests and providing additional experimental details about the re- analyzed datasets.

## Supplemental Figure Legends

**Figure S1. Bioinformatics pipeline to identify mouse-specific sequencing reads in scRNA-seq data.** Sequencing reads from the single-cell transcriptomes were deduplicated and aligned against the human genome using bowtie aligner with no mismatches allowed (v=0). Unaligned reads were aligned to mouse genome to reveal mouse-specific reads followed by deduplication, removal of low-complexity reads, and final precise filtering using the *blastn* algorithm. Both mouse- and human-specific reads were counted with HTSeq-count (*stranded=no* mode).

**Figure S2. Full-length mouse transcripts detected in nonLRC cells.** Alignment of mouse-specific sequencing reads to Pdcd10, Fth1, and Ftl1 mRNA transcripts from re-isolated nonLRC cells. Exons are represented by tall and short orange rectangles, indicating the coding and non-coding regions, respectively. Introns are depicted as lines with arrows pointing from the 5’ end to the 3’ end. Every lane corresponds to a single re-isolated human cell. The Y axis shows the number of reads.

**Figure S3. Sets of mouse transcripts detected in LRC (A) and nonLRC (B) human cells.** The number of unique mouse-specific reads for each transcript (y axis) detected in each cell (x axis) is reflected by the size of the black circle according to the legend. Large vertically aligned circles represent abundant mouse transcripts found in rare individual cells, likely indicating contamination. Frequent horizontal circles, both small and large, represent rare transcripts found in sets of human cells, suggesting their transfer into the human cells.

**Figure S4. Detection of Ftl1 transfer in *in vitro* co-culturing system with HeLa cells as acceptors.** A, panoramic view of the area occupied with seeded HeLa acceptor cells and donor RAW264.7 cells migrated towards them (maximum intensity projection). B, representative examples from A of Ftl1 transcripts localized within the areas occupied by the acceptor cells (marked with white arrows). Cyanine 3 was used to visualize FISH probes to Ftl1-specific padlock probe. 4’,6-diamidino-2-phenylindole (Dapi), was used to visualize genomic DNA.

