## Supplementary figures and images for "Evidence for *in vivo* mRNA Transport Between Mammalian Cells"

### Supplemental figure 1

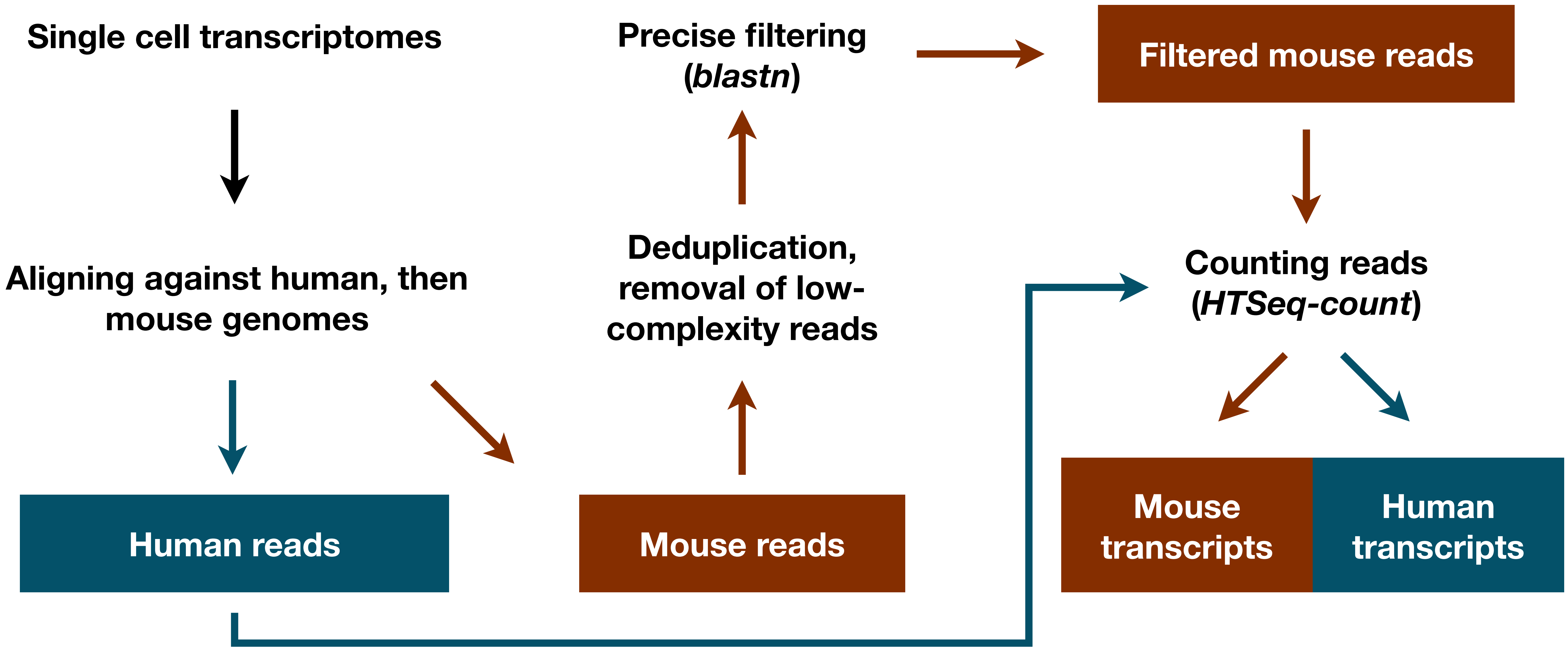

### Supplemental figure 2

#### Pdcd10

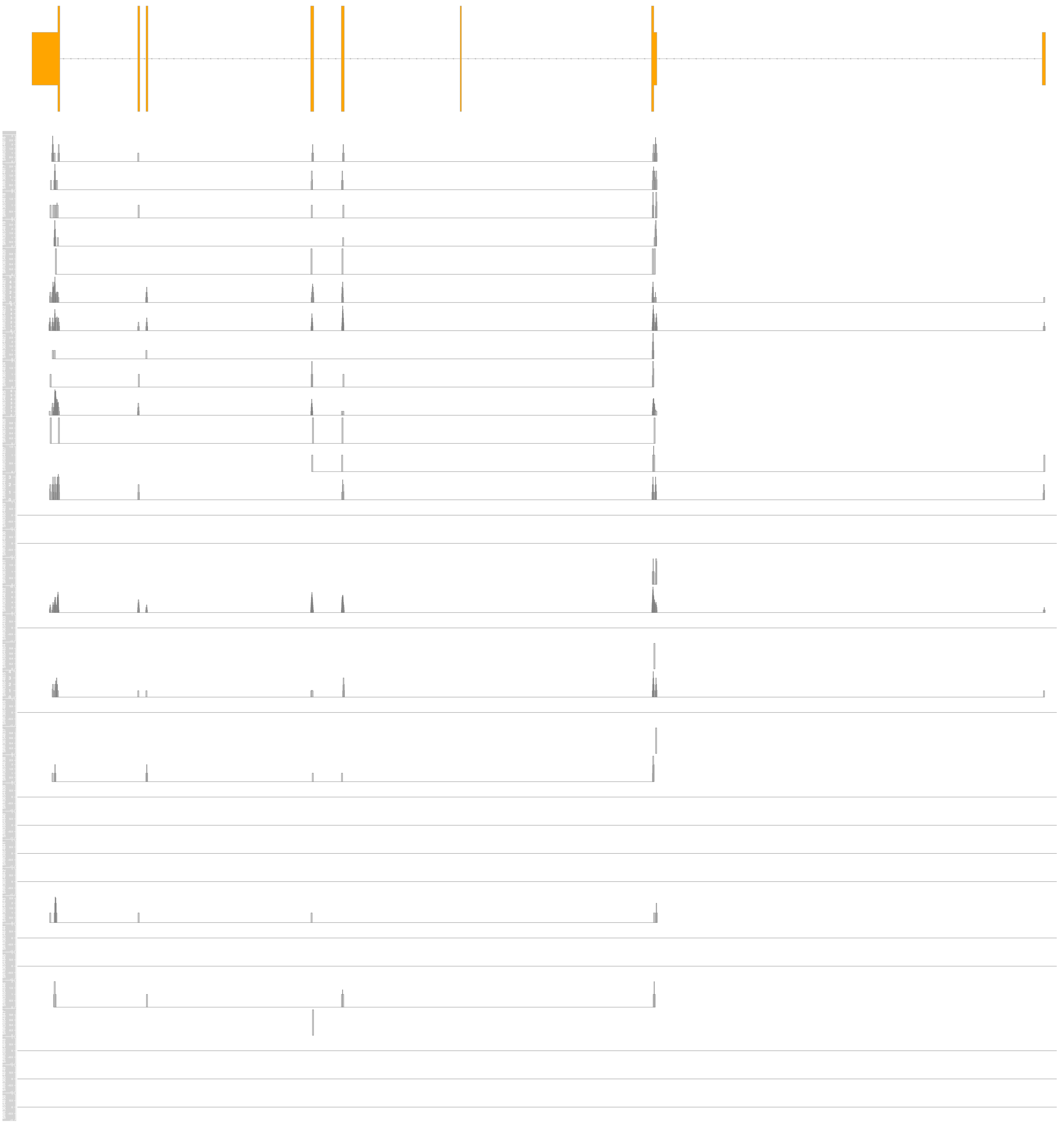

#### Fth1

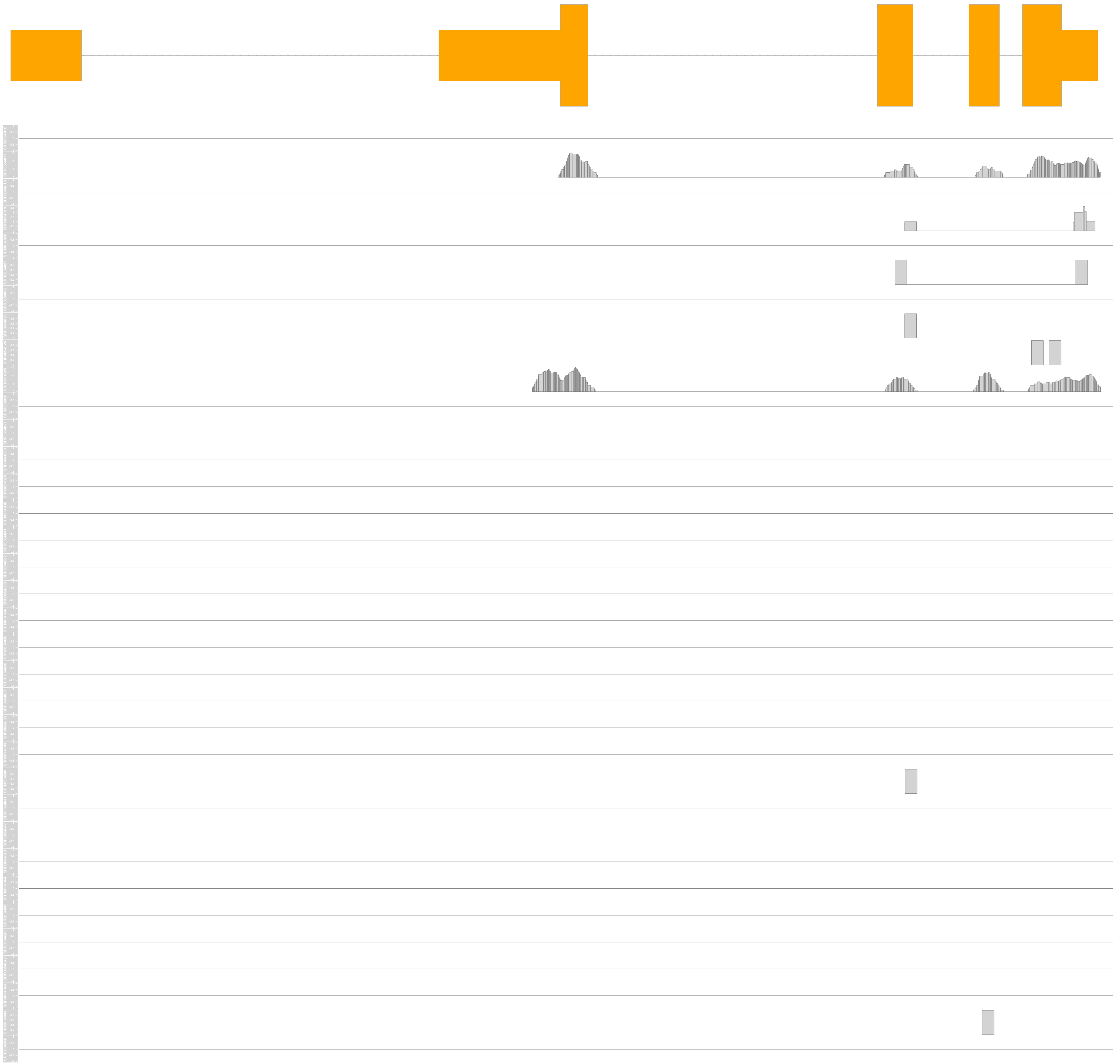

#### Ftl1

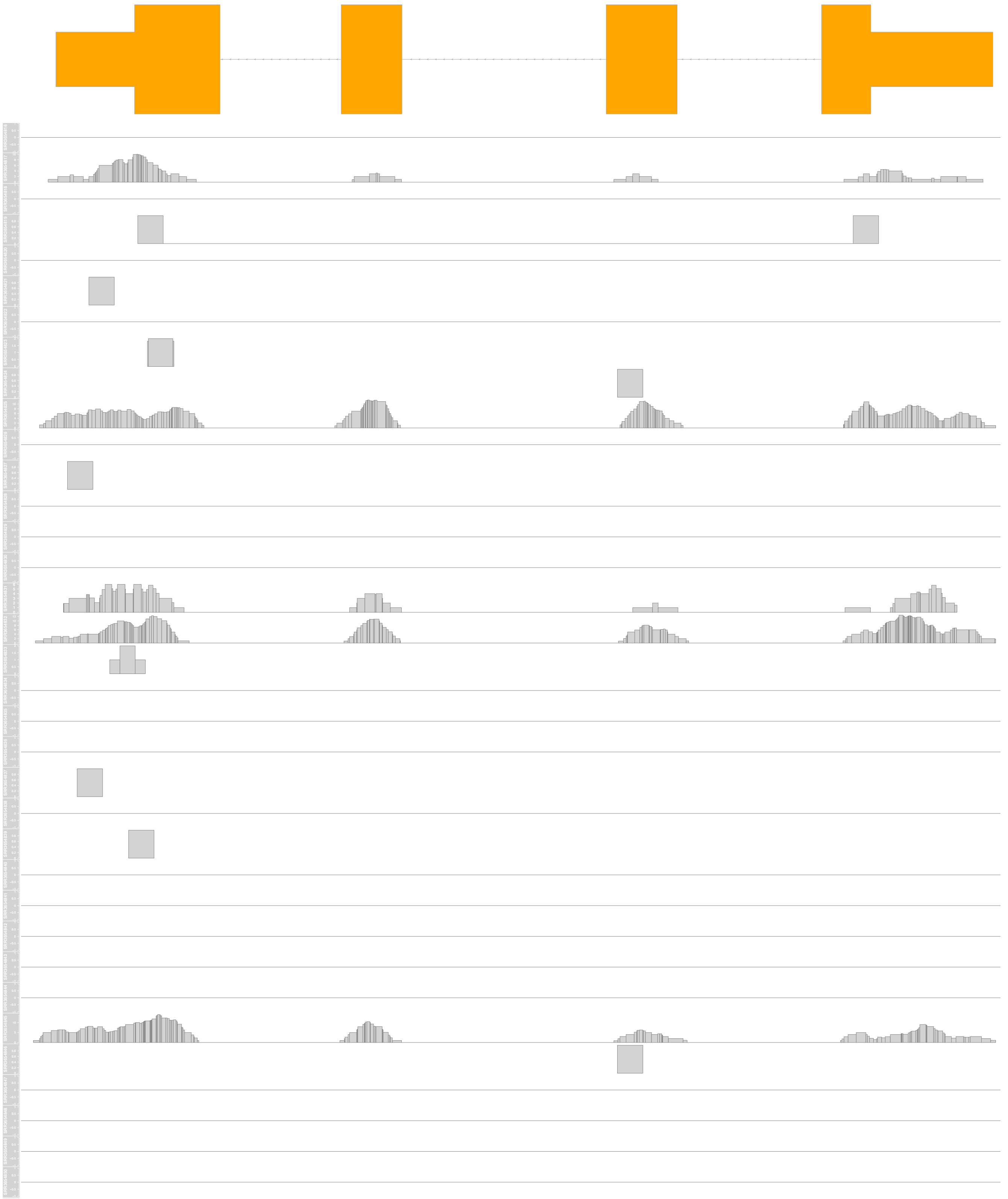

### Supplemental figure 3

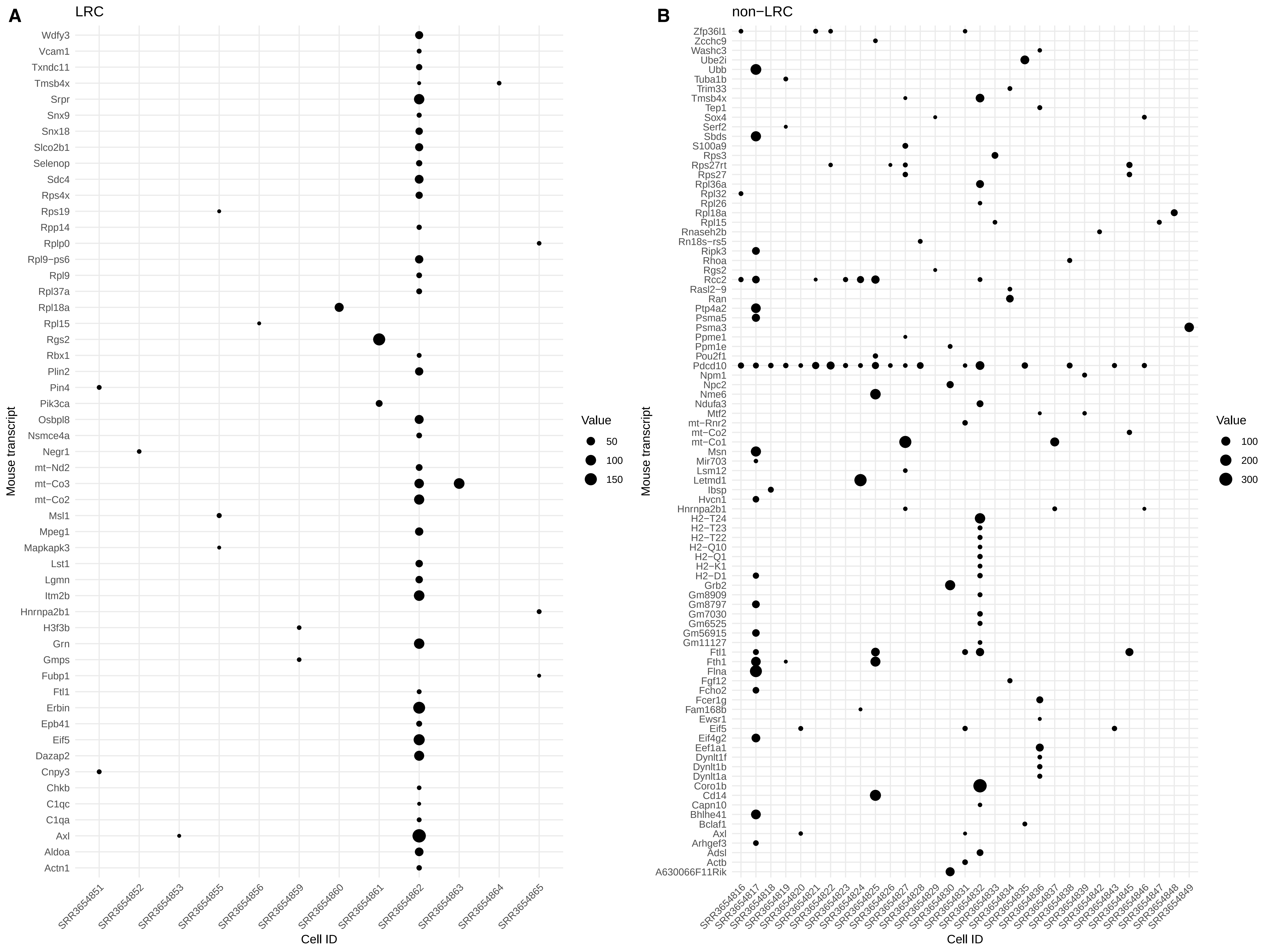

### Supplemental figure 4

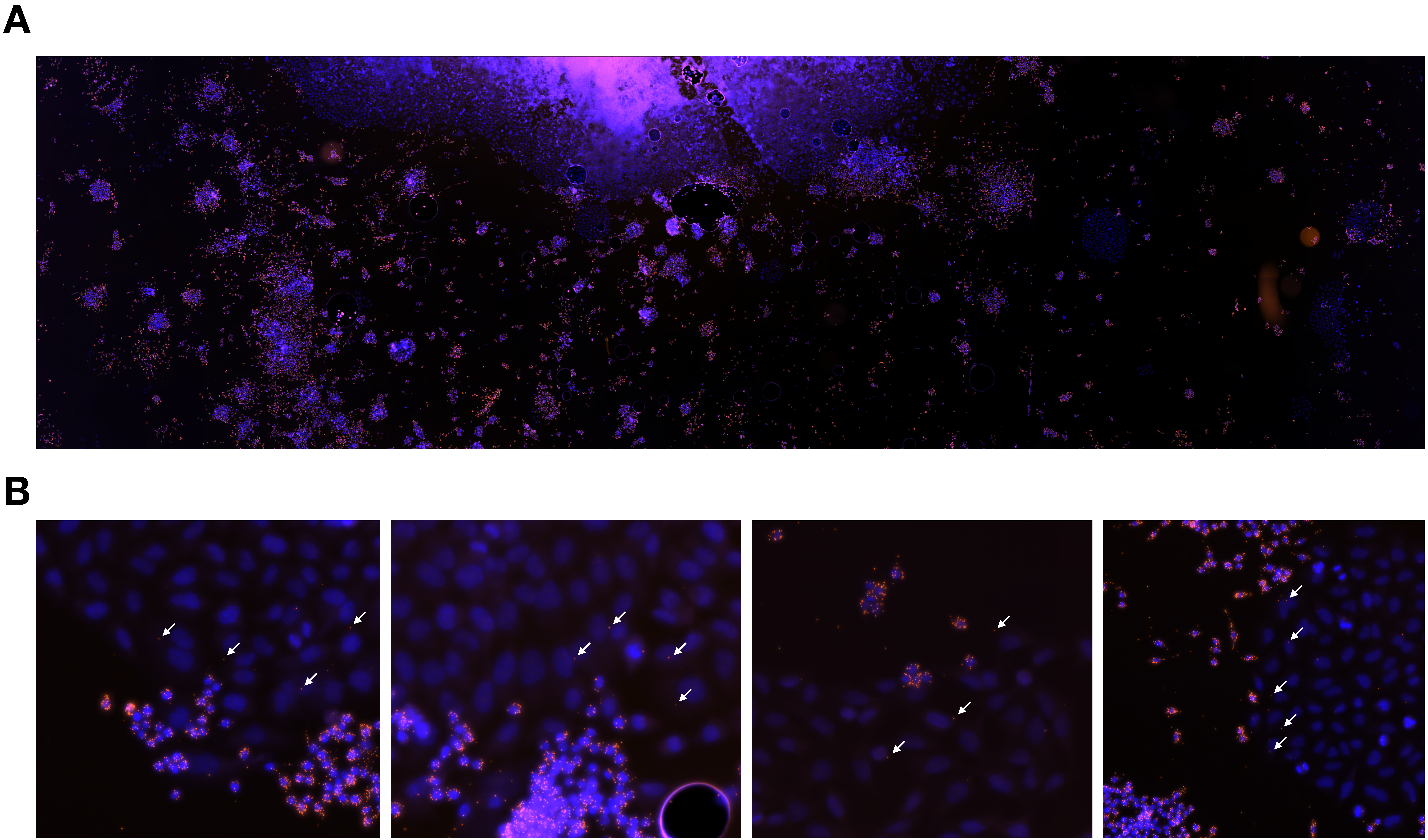
